# Assessment of mitochondrial physiology of murine white adipose tissue by mechanical permeabilization and lipid depletion

**DOI:** 10.1101/2020.02.19.956870

**Authors:** Natália C. Romeiro, Caroline M. Ferreira, Marcus F. Oliveira

## Abstract

White adipose tissue (WAT) is classically associated with energy storage in the form of triacylglycerol and is directly associated with metabolic disorders, including obesity. Mitochondria regulates cellular expenditure and are active in WAT. Although isolated mitochondria have been classically used to assess their functions, several artifacts can be introduced by this approach. Although methods to assess mitochondrial physiology in permeabilized WAT were proposed, important limitations that affect organelle function exist. Here, we established and validated a method for functional evaluation of mice mesenteric WAT (mWAT) mitochondria by using mechanical permeabilization in combination with lipid depletion and high-resolution respirometry. We observed that mild stirring of mWAT for 20 minutes at room temperature with 4% fatty acid-free albumin selectively permeabilized white adipocytes plasma membrane. In these conditions, mWAT mitochondria were intact and coupled, exhibiting succinate-induced respiratory rates that were sensitive to classical modulators of oxidative phosphorylation. Finally, the respiratory capacity of mWAT in females was significantly higher than in males, an observation that agrees with reported data using isolated mitochondria. The functional assessment of mWAT mitochondria through mild mechanical permeabilization, lipid depletion and high resolution respirometry proposed here will contribute to a better understanding of WAT biology in several pathophysiological contexts.

## 1. Introduction

White adipose tissue (WAT) represents the main energy store in the human body, responsible for the accumulation of high levels of triacylglycerides in large lipid droplets within white adipocytes. This enormous energy reserve is variable, depending not only on the energy demand, but also on the nutritional status of the organism [1]. Also, WAT is considered an important endocrine organ, responsible for secreting hormones such as adiponectin and leptin that regulate body’s energy metabolism and appetite, respectively [2]. Increased calorie intake, combined with a reduction in energy expenditure, shifts the energy balance towards the accumulation of triglycerides in WAT, which directly contributes to the onset of obesity and metabolic disorders [3]. This energy imbalance leads to hyperplasia and/or hypertrophy of adipose tissue [4,5, 6]. Therefore, strategies aiming the increase of energy expenditure in adipose tissues represent valid approaches to combat obesity and metabolic diseases.

Mitochondria are central organelles in cellular energy homeostasis and are known to be active in WAT, since mature adipocytes require ATP to maintain processes such as fatty acid activation and protein synthesis [7]. The routine methods for assessment of mitochondrial function are usually applied to whole intact cells, permeabilized tissue/cells and isolated mitochondria. Although most of the studies devoted to evaluate mitochondrial function used isolated organelles, important limitations generated by this approach must be considered, including the large sample effort, potential selection of mitochondrial subpopulations [8], and changes in mitochondrial morphology, also altering the organelle function [9]. For this reason, the use of permeabilized cells and tissues have been applied to overcome these issues, and provide advantages such as the possibility of evaluating mitochondrial physiology while preserving the structure and associations with other organelles [10].

The knowledge on mitochondrial function in WAT was essentially obtained by using isolated organelles from tissue biopsies. Recently, an efficient method to assess respiration in brown adipose tissue (BAT) biopsies was described [11], and was based on chemical permeabilization and depletion of tissue free fatty acids. Although alternative approaches to study WAT mitochondria in tissue biopsies were developed [12,13, 14], important limitations regarding the organelle integrity still remain. Critically, the use of detergents to chemically permeabilize plasma membrane is challenging and may affect the structure and integrity of mitochondrial membranes [15,16]. Importantly, given the huge lipid depots in white adipocytes, and the toxic effects of free fatty acids on mitochondrial structure and function [17, 18, 19], proper removal of these species is required [11, 20]. Thus, methods for studying mitochondrial function in WAT that preserve the different mitochondrial subpopulations, as well as their interactions with other organelles, are highly appreciated. Mature white adipocytes have an enormous lipid droplet that poses strong physical stresses on plasma membrane, rendering these cells sensitive to ruptures mediated by physical/chemical stimulus. Therefore, the present work proposes the development of a new method to evaluate mitochondrial physiology in mice mesenteric WAT (mWAT) biopsies. This new method allows the assessment of mitochondrial function in tissue biopsies and can be used to better understand WAT biology and the mechanisms that regulate the fat accumulation in this tissue.

## 2. Material and methods

### 2.1. Ethics statement

This research followed the ethics of animal use in research with a protocol approved by the Institutional Committee for the Care and Use of Animals at the Federal University of Rio de Janeiro (CONCEA 01200.001568/2013-87, protocol #016/19), and also followed the National Institutes of Health guide for the care and use of Laboratory animals (NIH Publications No. 8023, revised 1978).

### 2.2. Animals

All animals were fed an *ad libitum* chow diet and maintained at 19 - 22 ° C with a 14:10 h light-dark cycle until euthanasia performed by asphyxiation in a CO_2_ atmosphere and subsequent cardiac puncture. Mesenteric white adipose tissue (mWAT) from 4-month-old male and female C57BL/6J mice was carefully dissected in BIOPS buffer without ATP (10 mM Ca-EGTA; 0.1 µM calcium; 20 mM imidazole; 20 mM taurine; 50 mM K-MES; 0.5 mM DTT; 6.56 mM MgCl_2_; pH 7.1). Special care was taken to remove all visible blood vessels, tissues beyond WAT and fur.

### 2.3. Permeabilization methods

Dissected mWAT was weighted and subjected to different permeabilization methods: For mechanical + chemical permeabilization, the tissue was cut into small pieces in BIOPS buffer without ATP (10 mM Ca-EGTA; 0.1 µM calcium; 20 mM imidazole; 20 mM taurine; 50 mM K-MES; 0.5 mM DTT; 6.56 mM MgCl_2_; pH 7.1) and subjected to magnetic stirring (IKA^®^C-MAG MS 4, Brazil) of 100 rpm for 20 minutes at room temperature in BIOPS + 0.1 % (w/v) of free fatty acid albumin (FAF-BSA). Then, the solution was replaced with 10 mL of BIOPS + 0.05 mg/mL of saponin and the tissues were subjected to further 50 minutes of magnetic stirring of 100 rpm at room temperature. Then, the tissues were weighed and 9 mg biopsies were transferred to O2k respirometer chambers (Oroboros, Innsbruck, Austria) containing 2 mL of an adapted MIR05 buffer (0.5 mM EGTA; 3 mM MgCl_2_; 60 mM K-MES; 20 mM taurine; 10 mM KH_2_PO_4_; 20 mM HEPES; 110 mM sucrose; 0.5 % FAF-BSA; pH 7.2).

For mechanical permeabilization, dissected and weighted mWAT was cut into small pieces in BIOPS buffer without ATP and subjected to magnetic stirring of 100 rpm for 20 minutes at room temperature in BIOPS + 0.1 % (w/v), 4 % or 8 % FAF-BSA. Then, the solution was replaced by 10 mL of BIOPS without FAF-BSA and the tissues were subjected to further 50 minutes of stirring of 100 rpm at room temperature. Then, 9 mg biopsies of mWAT were weighed and transferred to O2k respirometer chambers (Oroboros, Innsbruck, Austria) containing 2 mL of modified MIR05 buffer using 0.5 % (permeabilized with 0.1 %), 4 % (permeabilized with 4 %) or 8 % (permeabilized with 8 %) FAF-BSA.

### 2.4. LDH activity

LDH activity was determined based on a protocol available in the literature [21]. An aliquot corresponding to 965 µL of each sample media were used: i) medium from tissues that have not underwent any process of permeabilization; ii) medium from tissues subjected to magnetic stirring in BIOPS buffer + FAF-BSA for 20 minutes; iii) medium from tissues subjected to magnetic stirring in BIOPS buffer only, for 50 minutes; iv) medium from tissues subjected to magnetic stirring in BIOPS buffer + 0.05 mg/mL saponin for 50 minutes, and v) medium from tissues which were homogenized in a Teflon-glass tissue grinder. Then, 0.3 mM NADH and 10 mM pyruvate were added to a quartz cuvette and the absorbance at 340 nm was recorded on a Shimadzu UV-2550 spectrophotometer (Shimadzu Scientific Instruments, Japan). Subsequently, the cuvette was incubated for 5 minutes at 37°C, and absorbance was registered again. The consumption of NADH was calculated based on the absorbance difference before and after incubation, using the 6.22 mM molar extinction coefficient of NADH at 340 nm. This assay was performed for the group that underwent permeabilization with 0.1 % and with 4 % FAF-BSA.

### 2.5. Respiration assessment

Cellular respiration was measured at 37°C and stirring at 300 rpm on a high resolution respirometer (Oroboros, Innsbruck, Austria) [15]. Permeabilized mWAT biopsies (9 mg) were added to 2 mL of modified MIR05 buffer (0.5 mM EGTA; 3 mM MgCl_2_; 60 mM K-MES; 20 mM taurine; 10 mM KH_2_PO_4_; 20 mM HEPES; 110 mM sucrose; 0.5 % (permeabilized with 0.1 %), 4 % (permeabilized with 4 %) or 8 % (permeabilized with 8 %) FAF-BSA; pH 7.2). For the evaluation of respirometry, the following routine was used: Initially, a volume of O_2_ gas was injected to a final concentration of 400 nmol/mL in the chamber. Then, 1.25 µM rotenone was added to prevent electron flow from succinate dehydrogenase to complex I (electron backflow). Succinate (10 mM final concentration) was added as a substrate to sustain respiration and also worked as quality control to assess the efficiency of tissue permeabilization, as succinate transport in plasma membrane of most eukaryotic cells is negligible [22]. Thus, activation of respiratory rates upon succinate addition indicates the efficiency of the permeabilization. ATP-linked respiration as a result of oxidative phosphorylation (OXPHOS) was assessed by titrating ADP to a final concentration of 4 mM. The maximum respiratory capacity was promoted titrating the proton ionophore FCCP up to 6 µM. Complex III was subsequently inhibited by adding 1 µM antimycin A (AA). As a quality control test for the integrity of the mitochondrial outer membrane [23], 10 μM cytochrome c was added to each respirometer chamber. In this context, the absence of a significant stimulating effect (<10%) on respiration indicates minimal damage of outer mitochondrial membrane during permeabilization. As a quality control of the selectivity of permeabilization of plasma membrane, four OXPHOS modulators were tested. Complex III inhibition by AA (1 µM) was added upon FCCP; Complex II inhibition by malonate (15 mM) upon FCCP; The coupling between respiration and mitochondrial ATP synthesis of mWAT was determined in two different ways: i) by carboxyatractyloside (CAT, 100 µM) treatment upon ADP, as CAT blocks the adenine nucleotide translocator activity; ii) oligomycin (2.5 µM) treatment upon ADP, as oligomycin selectively inhibits F1Fo ATP synthase. The drops in respiratory rates by using these inhibitors indicate the degree of functionality or coupling of the mitochondria.

### 2.6. Cytochrome c oxidase (COX) activity

The mWAT (9 mg) was homogenized in hypotonic buffer (25 mM K_2_PO_4_; 5 mM MgCl_2_; pH 7.4) in a glass and Teflon homogenizer on ice [24]. The suspension was centrifuged at 300 x g for 30 seconds and the supernatant was used to quantify protein levels [25]. An aliquot corresponding to 60 µg of protein was added to a cuvette containing 1 mL MIR05 buffer, 50 µM reduced cytochrome c and 1 µM AA, and the absorbance was measured at 550 nm in a Shimadzu UV-2550 spectrophotometer (Shimadzu Scientific Instruments, Tokyo, Japan) at 37°C. After 1 minute of reaction, 1 mM KCN was added to the chamber to inhibit all COX activity. This activity was calculated based on the difference in the oxidation rate of cytochrome c before and after the addition of KCN, according to the molar extinction coefficient of cytochrome c (18.7 mM). Activity was expressed as enzyme units (µmol of product/minute/mg of protein). COX activity was also determined by an independent method. Briefly, 2 mM ascorbate and 0.5 mM TMPD were added as an electron-donor regenerating system to reaction media (MIR05) containing 2.5 μg/mL AA. To distinguish cellular respiration from TMPD chemical auto-oxidation, 5 mM KCN was added at the end of each experiment, and COX activity was considered as the KCN-sensitive rate of oxygen consumption. Activation of respiration by the TMPD+ascorbate system was also used as a read-out of permeabilization efficiency, once TMPD is not permeable to intact plasma membrane.

### 2.7. Statistical analyses

Statistical analyses were carried out in the GraphPad Prism software version 6.00 for Windows (GraphPad Software, USA). D’Agostino and Pearson normality tests were done for all values to assess their Gaussian distribution. To compare experiments between three or more groups, we used the non-parametric Kruskal-Wallis test. When significant effects of groups or interaction between groups and states were obtained, we applied the Dunn’s Multiple Comparison post-test. For pair wise comparisons Student’s t tests (when there was a normal distribution) or the Mann-Whitney test (nonparametric) were performed. Statistical significance was established for p<0.05. All data were expressed as mean ± standard deviation.

## 3. Results and discussion

### 3.1. Establishment of permeabilization methods

In this paper, we designed a method to study mitochondrial physiology *in situ* using mechanically permeabilized mesenteric white adipose tissue (mWAT) from adult mice. To assess mitochondrial O_2_ consumption in mWAT biopsies, we subjected the samples to different permeabilization procedures, which was based on a similar method developed for mice BAT [11]. In comparison to WAT, brown adipocytes accumulate less lipids, releasing less free fatty acids during permeabilization and thus less prone to physical stresses and cell lysis. For these reasons, we made several adaptations of the original protocol developed for BAT, to assess mitochondrial physiology in mWAT and the experiments were performed as shown in Figure 1.

**Figure 1.**
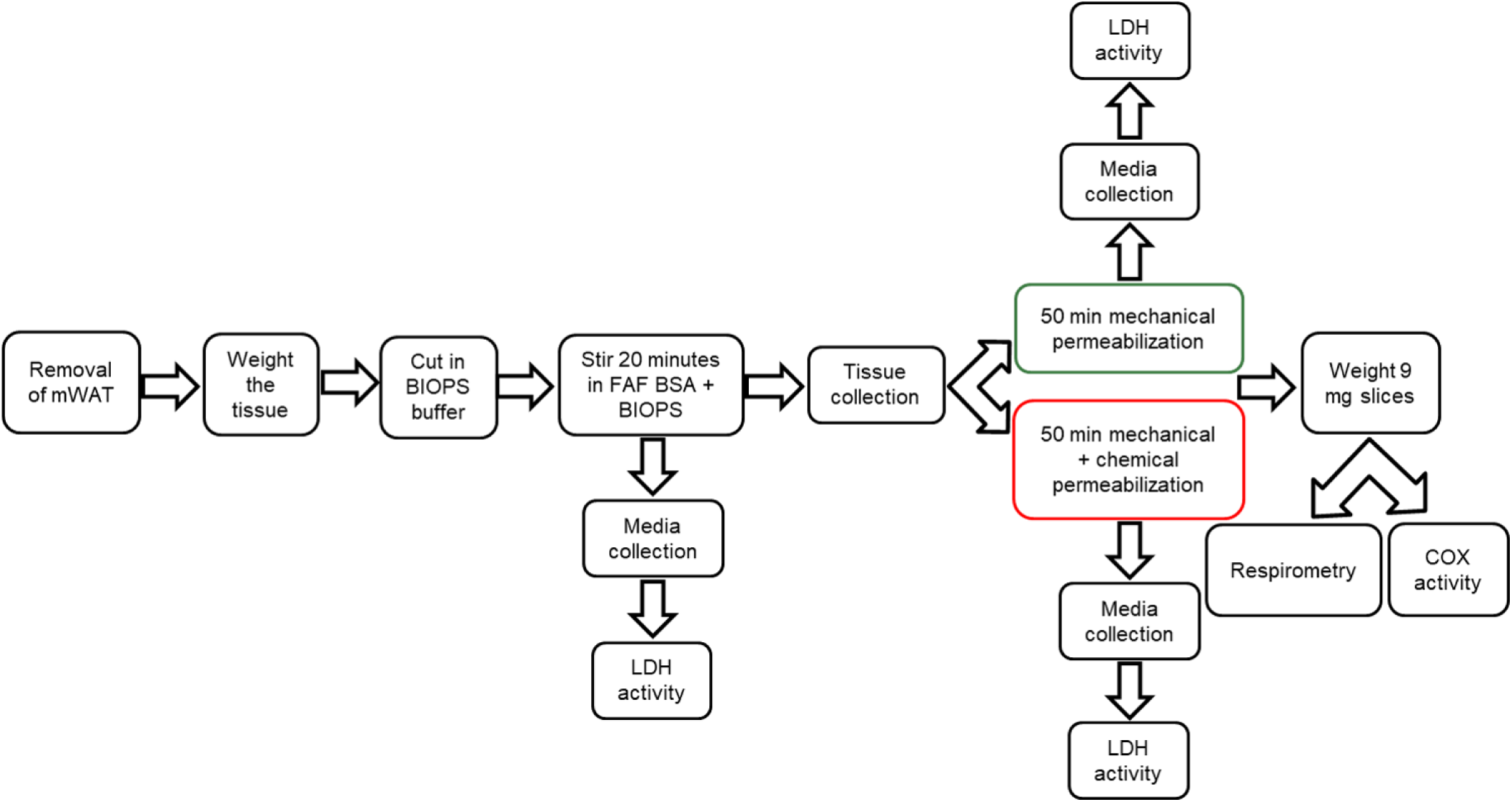
Schematic representation of the experimental approach used in this work to assess mWAT mitochondrial physiology by mechanical permeabilization and lipid depletion. mWAT was dissected from C57BL/6J mice, weighed, cut into small pieces and subjected to 20 minutes of magnetic stirring in presence of BIOPS + FAF-BSA, when the medium was collected for LDH activity. Then, tissue biopsies were transferred to BIOPS + saponin and subjected to further 50 minutes of stirring (mechanical + chemical permeabilization) or to BIOPS only under 50 minutes of stirring (mechanical permeabilization). After this step, the medium was collected for LDH activity and the tissues were weighed for biochemical analyses.

### 3.2. Mild magnetic stirring is sufficient to permeabilize white adipocyte plasma membrane, but proper functional assessment of mitochondria requires optimal lipid depletion by FAF-BSA

Our goal was to assess mitochondrial functionality of mWAT by selective permeabilization of the adipocyte plasma membrane. Firstly, we evaluated the activity of a cytosolic marker enzyme in the medium upon the permeabilization steps. Lactate dehydrogenase (LDH) is essentially a cytosolic enzyme that performs the redox interconversion of pyruvate and lactate using NADH and NAD+ as electron transfers. LDH activity was then determined in media from mWAT biopsies that i) were not permeabilized; ii) remained for only 20 minutes under magnetic stirring in BIOPS + 0.1 % FAF-BSA solution; iii) was subjected to magnetic stirring for 50 minutes in BIOPS; iv) was subjected to magnetic stirring for 50 minutes in BIOPS + 0.05 mg/mL saponin and v) was previously homogenized in a tissue grinder. Figure 2A shows that LDH activity of media from mWAT subjected to either mechanical or mechanical+chemical permeabillization was roughly identical to media from homogenized tissue. Importantly, LDH activities from these groups were significantly higher from media of non-permeabilized mWAT. Thus, we conclude that both types of approaches were capable to disrupt white adipocyte plasma membrane. Our next step was to evaluate mitochondrial functionality of permeabilized mWAT using both methods. The permeabilization procedure for BAT, which we based our method [11], used 0.1 % FAF-BSA to deplete free fatty acids and triglycerides, an important step to avoid lipotoxicity and uncoupling respiration from ATP synthesis [26]. However, mWAT respiratory rates under 0.1 % FAF-BSA were somewhat low, very noisy and loosely coupled in mWAT (Figure 2B). mWAT oxygen consumption rate (OCR) from groups subjected to mechanical or mechanical + chemical permeabilization were essentially the same, regardless the metabolic condition (proton leak-associated respiration, Figure 2C; ATP-linked respiration, Figure 2D). Indeed, the sub-optimal conditions under 0.1 % FAF-BSA were clearly seen upon addition of the proton ionophore FCCP, which unexpectedly reduced respiratory rates. This clearly indicates that 0.1 % FAF-BSA was not enough for adequate lipid depletion in permeabilized mWAT, resulting in accumulation of free fatty acids and potentially generating artifacts (Figure 2E). Assessment of cytochrome c oxidase (COX) activity, through TMPD + ascorbate induced oxygen consumption, can only be performed in cells where the plasma membrane was permeabilized [27]. Therefore, to assess mWAT plasma membrane permeabilization, we compared COX activity from tissue biopsies submitted to 30 and 100 minutes of magnetic stirring in the respirometer chamber. Figure 2F shows that COX activity was very low in mWAT that were stirred for 30 minutes, compared to samples kept for 100 minutes. We concluded that magnetic stirring of mWAT for 100 minutes permeabilizes white adipocyte plasma membranes (Figure 2A). However, poor lipid depletion by 0.1 % FAF-BSA prevented the accurate assessment of mitochondrial physiology (Figures 2C-F).

**Figure 2.**
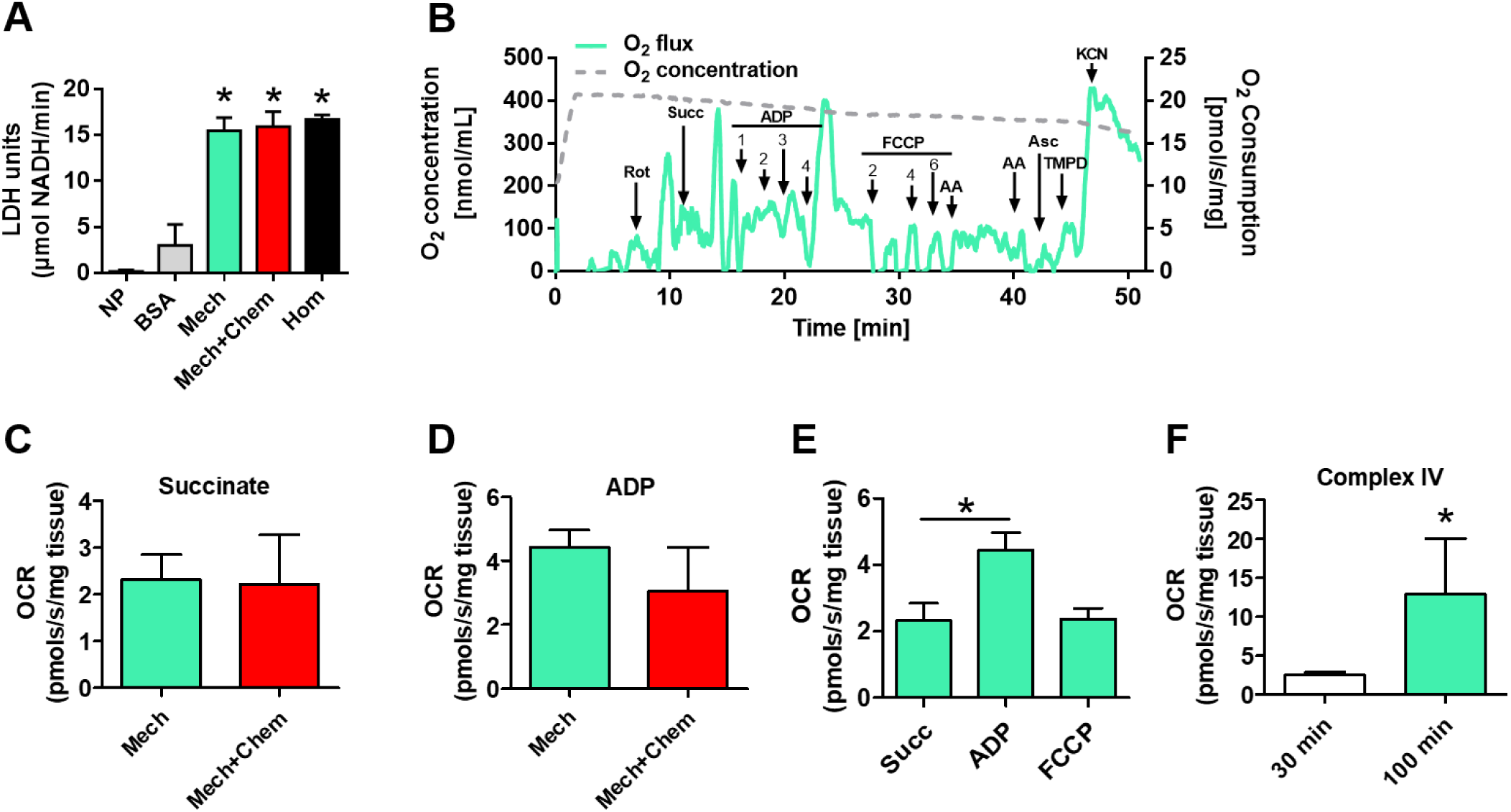
Magnetic stirring allows plasma membrane permeabilization of mWAT adipocytes but limited triglyceride depletion by 0.1% FAF-BSA does not preserve mitochondrial function. mWAT was subjected to either mechanical (Mech, green bars and line) or mechanical + chemical (Mech+Chem, red bars) permeabilization protocol using 0.1 % FAF-BSA. (A) LDH activity assessed in media from non-permeabilized tissues (NP), mWAT maintained with BSA only (grey bar), mechanical (green bar), mechanical + chemical (red bar) permeabilization and mWAT homogenates (Hom, black bar). (B) Representative trace of mWAT oxygen tension (grey line) and OCR (green line) after mechanical permeabilization in the presence of 0.1 % FAF-BSA. Numbers represent concentrations of ADP (mM) and FCCP (µM). Oxygen consumption rates (OCR) of mWAT under succinate (C) and succinate+ADP (D) that were subjected to either mechanical (green bars) or mechanical + chemical (red bars) permeabilization in the presence of 0.1 % FAF-BSA. (E) Quantification of mitochondrial respiratory states of mechanically permeabilized mWAT under succinate, ADP and FCCP treatments. (F) Cytochrome *c* oxidase (COX) activity mWAT subjected to mechanical permeabilization for 30 and 100 minutes. Data are expressed as mean ± standard deviation for at least three different experiments. Statistical comparisons between groups were carried out by using the Kruskal-Wallis and *post hoc* Dunn’s multiple comparison tests (A, E) or Mann-Whitney (C, D, F) test. * p<0.05, ** p<0.01.

### 3.3. The use of 4 % FAF-BSA during all steps makes respiration more coupled and maintains the integrity of the mitochondrial outer membrane

Previous studies have designed methods to assess mitochondrial physiology in WAT biopsies [11, 12, 13, 14]. However, the large triglyceride content, the potential release of these lipids during assessment of mitochondrial physiology, and the known cytotoxic effects of free fatty acids pose major challenges to determine organelle function in WAT [11, 17, 18, 19, 20]. In this sense, two critical technical aspects must be considered: i) proper assessment of mitochondria in white adipocytes through permeabilization of plasma membrane; ii) proper and continuous removal of excess triglyceride and other lipids during measurements. In this sense, the use of detergents and insufficient free fatty acid removal during respirometry experiments are important limitations [12, 13, 14]. We then reasoned that increasing FAF-BSA concentrations in permeabilization and respiration media would remove more efficiently excessive lipids, ultimately allowing optimal assessment of mWAT mitochondrial function. We firstly checked whether 4 % FAF-BSA would alter mWAT mechanical permeabilization. We then measured LDH activity from i) media of non-permeabilized mWAT; ii) media from mWAT kept for only 20 minutes under magnetic stirring in a BIOPS + 4 % FAF-BSA solution; iii) media from mWAT kept for 50 minutes under magnetic stirring only in BIOPS; iv) media from mWAT kept for 50 minutes under magnetic stirring in BIOPS + 0.05 mg/mL saponin and v) media from mWAT homogenized by a tissue grinder. Figure 3A shows that LDH activity in the media from mechanically permeabilized mWAT was very similar to either mechanical + chemical permeabilization and homogenization. Importantly, LDH activities of these three groups were significantly higher than non-permeabilized mWAT, indicating that 4 % FAF-BSA did not interfere with our procedure of white adipocyte plasma membrane permeabilization (Figure 3A). We next assessed the respiratory capacity of mWAT upon mechanical permeabilization in the presence of 4 % FAF-BSA. We observed that respiratory rates under 4 % FAF-BSA (Figures 3B and 3E) were substantially higher, more stable and more coupled than 0.1 % FAF-BSA (Figure 2E). This strongly indicates that the low quality of the results obtained with 0.1 % FAF-BSA were indeed artifacts generated by the excess of free lipids. Interestingly, the respiratory profile of mWAT mechanically permeabilized under 8 % FAF-BSA revealed to be inadequate, strongly reducing respiratory rates after ADP and FCCP additions (Figure 3E). Therefore, 4 % FAF-BSA provides the optimal condition for assessment of mWAT respiratory capacity.

**Figure 3.**
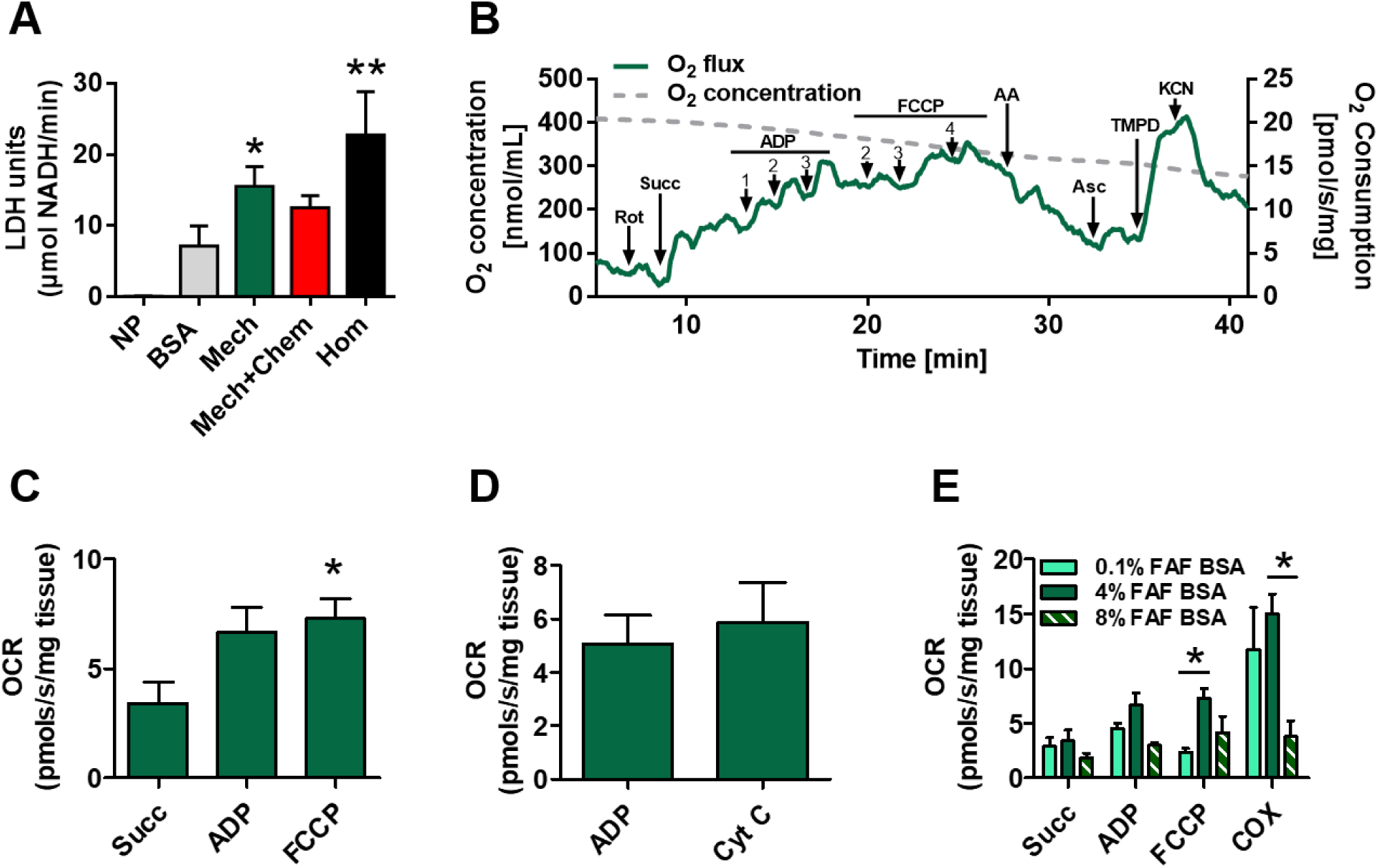
Mechanical permeabilization of mWAT in the presence of 4% FAF-BSA provides the optimal condition for assessing mitochondrial function. mWAT was subjected to either mechanical (Mech, dark green bars and line) or mechanical + chemical (Mech+Chem, red bar) permeabilization protocol using 4 % FAF-BSA. (A) LDH activity assessed in media from non-permeabilized tissues (NP), mWAT maintained with BSA only (grey bar), mechanical (dark green bar), mechanical + chemical (red bar) permeabilization and mWAT homogenates (Hom, black bar). (B) Representative trace of mWAT oxygen tension (grey line) and OCR (dark green line) after mechanical permeabilization in the presence of 4 % FAF-BSA. Numbers represent concentrations of ADP (mM) and FCCP (μM). (C) Quantification of mitochondrial respiratory states of mechanically permeabilized mWAT under succinate, ADP and FCCP treatments. (D) Effect of 10 µM cytochrome *c* supplementation on mWAT OCR subjected to mechanical permeabilization. (E) Quantification of mitochondrial respiratory states and COX activity of mechanically permeabilized mWAT subjected to mechanical permeabilization in the presence of 0.1 % (light green bars), 4 % (dark green bars) and 8 % FAF-BSA (hatched green bars). Data are expressed as mean ± standard deviation for at least three different experiments. Statistical comparisons between groups were carried out by using the Kruskal-Wallis and *post hoc* Dunn’s multiple comparison tests (A, C and E) or Mann-Whitney test (D). * p<0.05, ** p<0.01.

The set of data presented to this point led us to the conclusion that our method of white adipocyte plasma membrane permeabilization by mechanical stirring and lipid depletion allows optimal assessment of mWAT mitochondrial function. However, one can argue that the integrity of subcellular structures, especially the mitochondrial outer membrane was compromised by our permeabilization procedures. For this sake, we determined the effect of 10 µM cytochrome *c* supplementation on ATP-linked respiratory rates. Since mitochondrial outer membrane is impermeable to cytochrome *c*, a substantial increase (>10%) in OCR upon its addition is suggestive of disrupted mitochondrial outer membrane. We observed that cytochrome *c* did not cause significant increases in the respiratory rates of mWAT, ensuring that our permeabilization procedure preserves the integrity of mitochondrial outer membrane (Figure 3D). These results indicate that the use of 4 % FAF-BSA and mechanical permeabilization provides the ideal conditions for measuring mitochondrial OCR in mWAT biopsies.

### 3.4. Oxidative phosphorylation inhibitors reduce succinate-induced oxygen consumption in mechanically permeabilized mWAT

In the next set of experiments, we checked whether respiratory rates sustained by succinate oxidation in mechanically permeabilized mWAT would have the expected responses to classical OXPHOS inhibitors. We observed that antimycin A (AA), the classic inhibitor of complex III, significantly reduced respiratory rates by 41% under uncoupled conditions (Figure 4A). The classical inhibitor of complex II malonate also strongly reduced (∼72%) uncoupled respiratory rates (Figure 4B). In order to assess the integrity of mitochondrial inner membrane and OXPHOS coupling, we next investigated the effects of carboxyatractyloside (CAT), which inhibits the adenine nucleotide translocator (ANT, Figure 4C) and oligomycin, which blocks F1Fo ATP synthase (Figure 4D). Both CAT and oligomycin are powerful inhibitors of mitochondrial ATP synthesis by targeting mechanisms directly involved in ATP production, not in the electron transport system. Therefore, impairment of respiration provided by limited mitochondrial ATP synthesis avoids proton translocation through F1Fo ATP synthase, increasing the protonmotive force and ultimately reducing electron flow and respiration. Figures 4C and 4D show that blockage of mitochondrial ATP synthesis significantly reduced (∼ 52 and 54 %, respectively) ATP-linked respiration in mechanically permeabilized mWAT. Thus, the method to evaluate mitochondrial physiology in mWAT proposed here allows the adequate assessment of respiratory rates, substrate oxidation and OXPHOS coupling without organelle isolation or detergent use.

**Figure 4.**
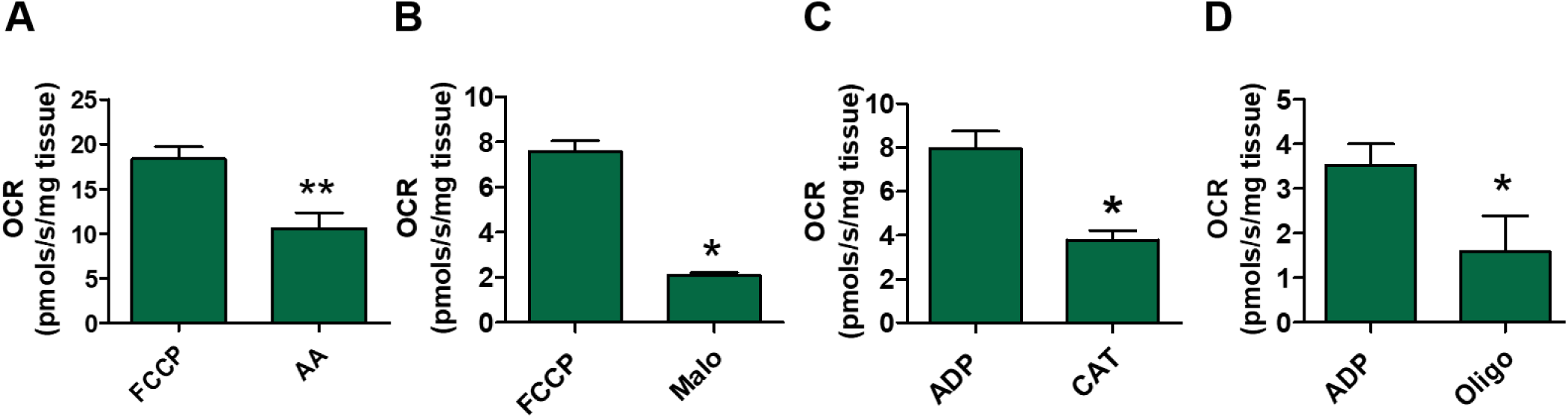
Oxidative phosphorylation inhibitors reduce succinate-induced respiratory rates in mechanically permeabilized mWAT. mWAT was subjected to the permeabilization protocol using 4 % FAF-BSA and tissue respiration was determined in the presence of 1.25 µM rotenone and 10 mM succinate. (A) Effect of inhibition of complex III by the addition of 1 µM AA in respiration uncoupled by 6 µM FCCP. (B) Effect of inhibition of complex II by the addition of 15 mM malonate on respiration uncoupled by 6 µM FCCP. (C) Effect of inhibition of the adenine nucleotide exchanger by adding 100 µM CAT to the coupled respiration after adding 4 mM ADP. (D) Effect of F1Fo ATP synthase inhibition by adding 2.5 µM oligomycin to the coupled respiration after adding 4 mM ADP. Data are expressed as mean ± standard deviation for at least three different experiments. Statistical comparisons between groups were carried out using the Student’s t (A) and Mann-Whitney (B, C, D) tests. * p<0.05, ** p<0.01.

### 3.5. Protocol validation of mWAT mechanical permeabilization by assessing sexual differences in respiratory rates

Evidence indicates that mitochondrial enzyme activities in WAT from women were reportedly higher than from men [28]. In order to validate the protocol described here, we assessed the respiratory capacity and COX activity from mWAT of male and female mice. We observed that females OCR were higher compared to males, regardless the respiratory state analyzed (Figure 5A). We also observed that COX activity, assessed by TMPD+ascorbate respirometry, was significantly higher in females than in males (Figure 5B). To ensure the reliability of our measurements, COX activity in both sexes was determined using an independent method (by following the oxidation of reduced cytochrome *c* at 550 nm). Indeed, we observed that COX activity was significantly higher in females mWAT compared to males (Figure 5C). Therefore, the method of mechanical permeabilization and lipid depletion for assessment of mitochondrial function in mWAT, proposed here, recapitulate the pattern observed from isolated mitochondria (Table 1) and validate this procedure for studies of energy metabolism in mice white adipocytes.

**Figure 5.**
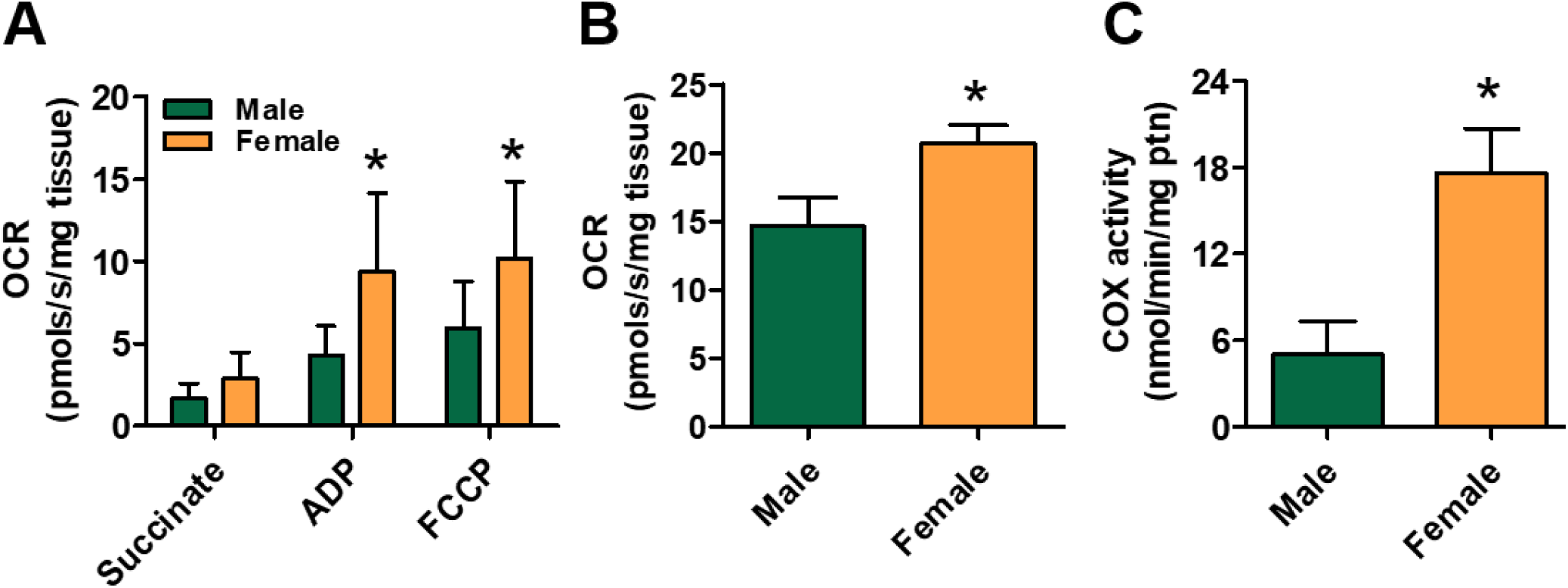
Method validation of mWAT mechanical permeabilization by assessing sexual differences in respiratory capacity. High resolution respirometry of mWAT from male (green bars) and female (orange bars) mechanically permeabilized mice in the presence of 4% FAF-BSA. (A) Biopsies of mWAT corresponding to 9 mg of tissue were added to the respirometer chamber containing 2 mL of respiration buffer + 4 % FAF-BSA. Measurements started after the addition of 1.25 µM rotenone and 10 mM succinate, followed by 2 mM ADP, 4 µM FCCP and 1 µM AA. (B) COX activity measured through oxygen consumption induced by TMPD + Asc and subtracted from OCR after adding 5 mM KCN. (C) COX activity measured by the reduced cytochrome *c* oxidation rate determined at 550 nm. Data are expressed as mean ± standard deviation for at least three different experiments. Statistical comparisons between groups were carried out using the Student’s t (B) and Mann-Whitney (A and C) tests. * p<0.05.

Despite methods to assess mitochondrial physiology in WAT biopsies were previously carried out [12-14], their ability to properly measure tissue/cell respiration are limited by several reasons. For example, the possibility that mixing/measures protocols of the extracellular flux analyzer could disrupt adipocyte plasma membrane, combined with insufficient lipid depletion during respirometry experiments, were not considered [12]. Indeed, this would explain why only ∼ 22 % of reported basal respiration was linked to ATP production by OXPHOS [12]. Although the integrity of adipocyte plasma membrane was not assessed, it is likely that reported oxygen consumption rates may have a mixed contribution of either “intact” and mechanically permeabilized cells, but without adequate mitochondrial substrates to sustain respiration [12]. Secondly, the absence of suitable lipid depletion system in the media to quench free fatty acids and other lipid species released upon adipocyte lysis, may generate artifacts in respirometry that prevent proper interpretation of the available data [11,12,17-20]. On the other hand, approaches for assessment of mitochondrial metabolism in WAT using chemical permeabilization successfully demonstrated respiration sustained by different substrates oxidation [13-14]. However, it is likely that absence of optimal lipid depletion *<u>during respirometry experiments</u>* have introduced potential artifacts on OCR. For example, respiration in isolated WAT mitochondria provided by ADP and FCCP were hardly increased relative to succinate alone, which may be explained by undesirable side effects of lipids present in the media [13-14]. This is especially critical when WAT biopsies from high fat diet animals were assessed, where respiratory rates were also not activated by either ADP or FCCP [13-14]. When using whole adipocytes from different WAT depots, reported respiratory rates obtained by high resolution respirometry (as in the present study) were essentially associated to proton leak rather than ATP production [13-14]. This raises the possibility that during respirometry experiments, continuous stirring of WAT biopsies, as shown in the present work (Figure 3), may have disrupted plasma membrane, resulting in lipids release in the media and ultimately promoting mitochondrial proton leak [13-14]. Conceivably, limited lipid depletion by low FAF-BSA concentrations is not enough to quench all lipids released in the media, providing sub-optimal conditions for proper assessment of mitochondrial function (Figure 2B and 2E). Indeed, as respirometry in the previous studies [12-14] and here (Figure 2) were carried out in media containing only 0.1 % FAF-BSA, clear artifacts were observed as pointed out above. Thus, we conclude that optimal lipid depletion <u>during all steps of tissue permeabilization and respirometry</u> is a critical issue to be considered when studying mitochondrial metabolism in WAT biopsies.

Although the present work significantly advances to study mitochondria in the context of WAT biology, some limitations must be considered in our methodology as, i) we conducted our studies using a single strain of laboratory mouse (C57Bl6/J), one WAT depot (mesenteric), and in young animals fed only on chow diet; ii) we only assessed mitochondrial metabolism in permeabilized cells, thus limiting the extrapolation of our data to intact white adipocytes and WAT; iii) since high FAF-BSA concentrations were used, the effects of hydrophobic drugs on respiratory rates might be a challenge; iv) also due to high concentrations of FAF-BSA in the media, we were unable to normalize respirometry data by protein levels. However, even with all these limitations, the methodology established here provides a suitable way to assess mitochondrial physiology in mice mWAT biopsies using mechanical permeabilization in combination with lipid depletion. Importantly, optimal lipid depletion is a necessary step not only during mechanical permeabilization but also during respirometry measurements.

## 4. Conclusions

Here, we tested and validated a new and simple method to assess mice mWAT mitochondrial physiology by mild mechanical permeabilization, lipid depletion with 4 % FAF-BSA and high resolution respirometry. This procedure by-passes classical time and labor extensive mitochondrial isolation methods and the use of detergents to permeabilize mWAT. This methodology represents not only a significant step for the study of WAT biology, but also a valuable tool to better understand the mechanisms that regulate mitochondrial function in white adipocytes and the potential consequences to metabolic diseases.

## Abbreviations

WAT: White adipose tissue
mWAT: Mesenteric white adipose tissue
BAT: Brown adipose tissue
FAF-BSA: Fat acid free albumin
LDH: Lactate dehydrogenase
OXPHOS: Oxidative phosphorylation
ADP: Adenosine diphosphate
ATP: Adenosine triphosphate
AA: Antimycin A
Succ: Succinate
Cyt c: Cytochrome c
CAT: Carboxyatractyloside
FCCP: Carbonyl cyanide 4-(trifluoromethoxy)phenylhydrazone
Oligo: Oligomycin
COX: Cytochrome c oxidase
KCN: Potassium cyanide
Asc: Ascorbate
TMPD: Tetramethyl-p-phenylenediamine
Rot: Rotenone
OCR: Oxygen consumption rate

## 6. Acknowledgments

We thank MSc Eduardo Ferreira and Prof. Antonio Galina for the kind support in the animal facility used in this work.

